# Molecular spikes: a gold standard for single-cell RNA counting

**DOI:** 10.1101/2021.07.10.451877

**Authors:** Christoph Ziegenhain, Gert-Jan Hendriks, Michael Hagemann-Jensen, Rickard Sandberg

**Author notes:** Correspondence to:* Rickard Sandberg. Contributed equally (Christoph Ziegenhain, Gert-Jan Hendriks).

## Abstract

Molecule counting is central to single-cell sequencing, yet no experimental strategy to evaluate counting performance exists. Here, we introduce *molecular spikes*, novel RNA spike-ins containing inbuilt unique molecular identifiers that we use to identify critical experimental and computational conditions for accurate RNA counting across single-cell RNA-sequencing methods. The molecular spikes are a new gold standard that can be widely used to validate RNA counting in single cells.

---

Single-cell RNA sequencing (scRNA-seq) is being widely used to dissect cellular states, types and trajectories^1^. Common to many single-cell technologies are counting strategies to mitigate the overcounting of amplicons derived from each RNA or DNA molecule. Typically, a random sequence, or unique molecular identifier (UMI), is added via the adapter oligos prior to DNA amplification and sequencing^2^, and this strategy has become standard for RNA counting in single cells^3–6^. Despite the widespread use of UMIs, no experimental strategies exist that can be used to systematically quality-control counting accuracy in new single-cell methods or variations in chemistries used. Furthermore, errors within the barcodes during amplification and sequencing necessitate subsequent *ad hoc* computational correction strategies. Several approaches for UMI error corrections to estimate RNA molecule counts have been proposed^7–9^, but so far there are no experimental ground-truth datasets enabling standardized benchmarking. Here, we developed novel mRNA spike-ins that carry a high diversity random sequence (i.e. an internal UMI) that we use to assess the RNA counting accuracy of popular scRNA-seq methods and computational correction strategies.

Randomized synthetic DNA sequences with minimal overlap to the human and mouse genomes were cloned into pUC19, together with a T7 promoter and a poly-A tail consisting of 30 adenine nucleotides (**Figure 1a**). Oligonucleotide libraries carrying 18 random nucleotides were inserted either into the 5’ or 3’ region of the synthetic sequence to construct the spike-UMI (spUMI) of the 5’ and 3’ molecular spike, respectively (**Figure 1b**). The resulting plasmid libraries were then used for in vitro transcription to produce molecular spike RNA pools (**Figure 1a**). To test the produced spikes, we added the 5’-molecular spike to single HEK293FT cells and prepared Smart-seq3 libraries^6^. The spUMIs from the molecular spike sequences were extracted from aligned reads and we similarly extracted the standard UMI sequence introduced on the Smart-seq3 template switching oligo. We verified that the 18 bp spUMI was indeed predominantly random (**Supplementary Figure 1a**). To counteract PCR and sequencing errors within the spUMIs on the molecular spikes, we investigated the appropriate error correction strategy. To this end, we calculated for each molecular spike spUMI the minimum edit distance (hamming distance) to the closest sequence within the cell and to 1000 randomly sampled molecular spike spUMIs from other cells. This analysis demonstrated that the 18bp spUMIs often showed an enrichment of spUMIs with one or two base errors within cells (**Supplementary Figure 1b**). Moreover, random sampling of sequences of 18bp length is unlikely to yield collisions in sequence space (∼68.7 billion sequences) at a hamming distance of 2. Therefore, we used a hamming distance of 2 to infer the exact number of molecular spike spUMIs present in each cell for the remainder of the experiments in this study, and we further excluded spUMIs that were over-represented across cells (**Methods**) to remove potential biases (**Supplementary Figure 1c**). By fitting an asymptotic non-linear model to the number of observed spUMIs sequences across cells, we estimated the complexity of the total 5’-molecular spike pool to 3.2 million, demonstrating that there were no unexpected bottlenecks in the cloning and production procedure (**Figure 1c**).

**Figure 1:**
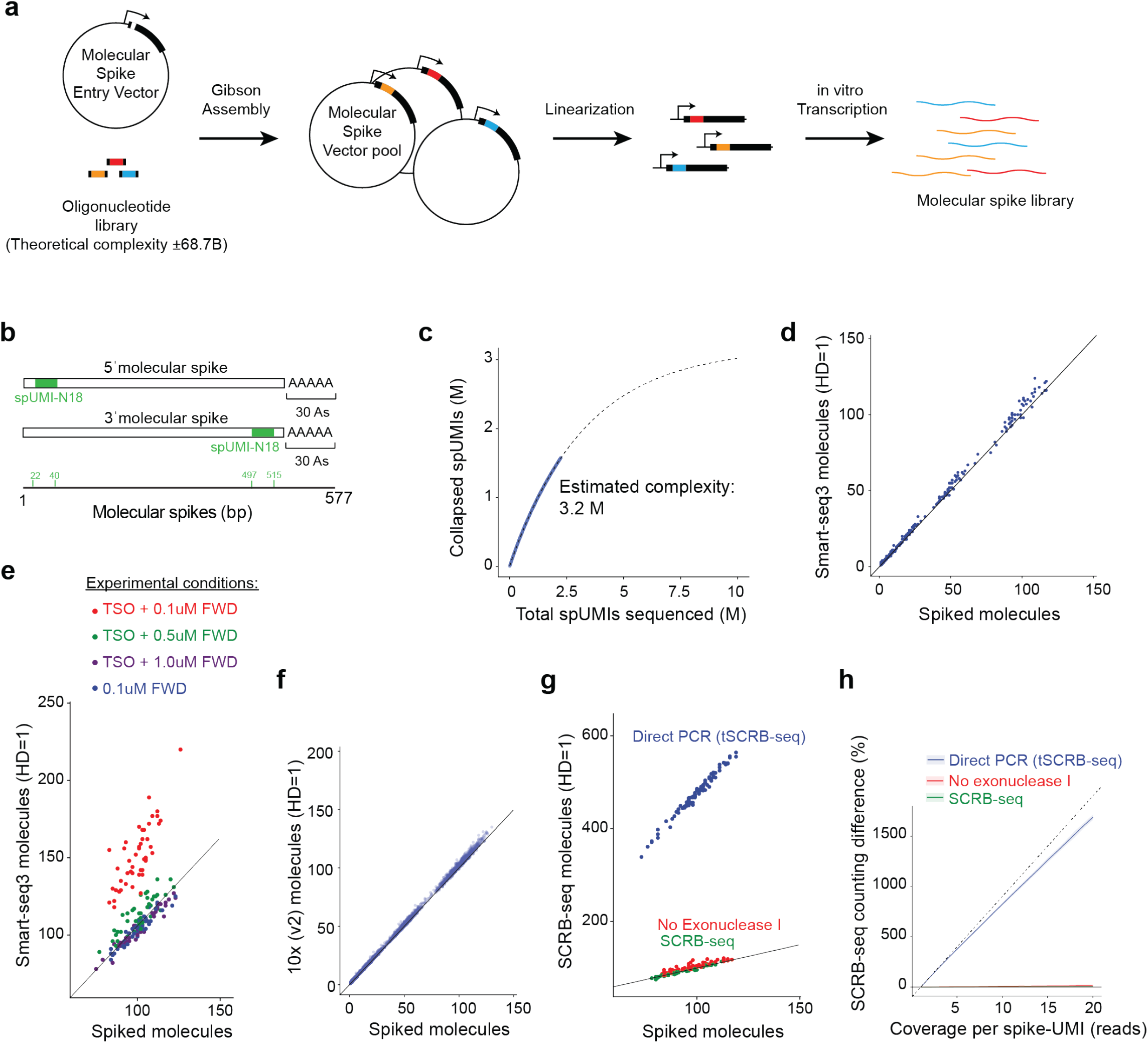
Direct assessment of single-cell RNA counting using molecular spikes. (**a**) Schematic of cloning strategy of molecular spikes, where an oligonucleotide library is inserted into a molecular spike entry vector, and the vector pool is linearized and in vitro transcribed to generate a complex molecular RNA spike-ins. (**b**) Coordinates of molecular spikes, with inbuilt UMI in the 5’ or 3’ end. (**c**) 5’-molecular spike complexity estimated by fitting a non-linear asymptotic model (dotted line) to unique spUMI sequences observed as a function of the number of spUMIs sequenced across cells (blue line). (**d**) Scatter plot showing error-corrected (hamming distance 1) Smart-seq3 RNA counts (y-axis) against the number of spiked molecules (x-axis) ranging from 1 to 100 spiked molecules per cell. Data from HEK293FT cells (n=48 cells). (**e**) Scatter plot showing number of spiked molecules (x-axis) against error-corrected RNA counts (hamming distance 1) for data generated with variations to the Smart-seq3 protocol, that utilize cDNA cleanup prior to amplification (0.1uM FWD) or without cleanup and therefore remaining TSO and for different concentrations of FWD primer. Data from 39 cells or more are shown per condition. (**f**) Scatter plot showing number of spiked molecules (x-axis) against error-corrected RNA counts (hamming distance 1) for 10x Genomics (v2) data (n=955 cells). (**g**) Scatter plot showing number of spiked molecules (x-axis) against error-corrected RNA counts (hamming distance 1) for data generated with variations to the SCRB-seq and tSCRB-seq protocols. Standard SCRB-seq (green, 53 cells), excluding Exonuclease I treatment (red, 77 cells) and direct PCR (tSCRB-seq) (blue, 90 cells). (**h**) Percent counting error (observed/true) for in RNA counts generated with variations to the SCRB-seq and tSCRB-seq protocols. Solid line denotes the mean over cells per condition with the shaded area representing the standard deviation colored by experimental conditions. Direct PCR (tSCRB-seq) (90 cells), No Exonuclease I (77 cells) and standard protocol (53 cells).

Having validated the randomness and complexity of the spUMI, we investigated the RNA counting accuracy of single-cell methods, starting with Smart-seq3^6^. Since the copy numbers of added 5’-molecular spikes were very high, we sampled molecular spike molecules from the range of expression levels typically found in HEK293FT cells (**Supplementary Figure 2**). The observed error-corrected Smart-seq3 counts closely followed (r^2^= 0.99) the molecular spike ground-truth (i.e. error-corrected spUMIs) (**Figure 1d**), demonstrating the accuracy in RNA counting in single-cells with Smart-seq3.

Next, we exemplify how the molecular spikes can properly diagnose inaccuracies in scRNA-seq library protocols by investigating altered Smart-seq3 conditions in which residual RNA-based template-switching oligo (TSO) is allowed to prime during PCR preamplification to cause artificially inflated RNA counts. Whereas the TSO can be efficiently removed by bead cleanups prior to PCR, it can also be effectively outcompeted by increasing concentrations of forward PCR primers (**Figure 1e**). However, the combination of remaining TSO with lower amounts of forward PCR primer results in significant TSO priming and inflation in RNA counting, at approximately 150% of the correct expression levels (**Figure 1e** and **Supplementary Figure 3**). We note that a minor count inflation (approximately 110%) is detectable even at 0.5 µM forward primer (at 100 RNA copies per cell and over 10 sequencing reads per molecule), and that an increase to 1.0 µM in Smart-seq3 effectively removes this remaining inflation.

Most scRNA-seq protocols rely on 3’-tagging mRNA instead of producing full-length coverage of transcripts, and we therefore engineered a 3’-molecular spike carrying the 18-nucleotide spUMI close to the poly-A tail (**Figure 1b**). After similar QC and filtering of 3’ spUMIs (**Supplementary Figure 4**), we first applied these molecular spikes to the droplet-generation process in 10x Genomics Gene Expression Assay (v2 chemistry; see **Methods**). The inferred molecule counts from this experiment were in good agreement with the molecular spikes (**Figure 1f**), as expected since the 10x Genomics protocol extensively purifies the cDNA prior to PCR amplification. Next, we applied the molecular spikes to the SCRB-seq protocol^10^, a plate-based 3’-tagging method that includes cDNA clean-up prior to cDNA amplification. The RNA counting in SCRB-seq was accurate (**Figure 1g**). Recently, tSCRB-seq^11^ was introduced and reported to have greatly increased sensitivity compared to SCRB-seq. In tSCRB-seq the PCR reagents are added directly to the individual reactions without cDNA clean-up. To assess how RNA counting was impacted in tSCRB-seq, we first generated a SCRB-seq library that omitted the Exonuclease I digest after reverse transcription, which is a safeguard against remaining oligo-dT primer potentially producing faulty amplicons in the subsequent PCR reaction, which resulted in minimal (105%) UMI counting inflation (**Figure 1g**). Following tSCRB-seq, we added PCR master mix directly into the individual wells of cDNA product and this “direct PCR” condition resulted in significant UMI overcounting (**Figure 1g**). In fact, the “direct PCR” implementation in tSCRB-seq introduced new UMIs nearly in every new sequenced read, resulting in overcounting that linearly follows sequencing depth (**Figure 1h**). Clearly, the UMI containing oligo-dT primer appears to be preferentially priming in the pre-amplification PCR reaction, introducing false new UMIs in every cycle. The clean-up after pooling RT products, even in the absence of the Exonuclease I digest, seemed to be very efficient at removing the oligo-dT primer. Thus, the reported increased sensitivity obtained in tSCRB-seq^11^ is completely artificial due to the removal of the essential cDNA cleanup step.

Having demonstrated the important role of molecular spikes in assessing the RNA counting abilities of scRNA-seq methods, we next systematically investigated UMI error-correction procedures and compared their inference to the ground-truth number of spiked in molecules. We based this analysis on the experiment with 10x Genomics using 3’-molecular spikes, and we sampled molecular spikes and their associated sequence reads (1 to 10 reads each) matching to 60 equally spaced expression levels between 1 and 1000 molecules (**Supplementary Figure 5a**). Moreover, we directly investigated the effect of the UMI length on the error-correction by performing these analyses in parallel on *in silico* trimmed versions of the observed 10-nucleotide 10x Genomics UMI. Basing the RNA counts on uncorrected UMI observations inflated the counts with increasing inflation in longer UMIs (**Figure 2a, b**) reflecting that longer UMI sequences have higher risks to be affected by PCR and sequencing errors. As expected, the inflated counts increased also with increasing read coverage and expression levels (**Supplementary Figure 5b**). Reassuringly, applying UMI error corrections that collapse UMI observations within a hamming distance of 1 (as implemented in the zUMIs pipeline^8^) removed a large proportion of counting errors for the longer UMI lengths (**Figure 2c,d**) and fully removed the dependency on coverage (**Supplementary Figure 5c**). In contrast to a previous report^12^, we observe that UMIs of a length 6 or lower reach significant collision rates leading to under-counting even in the absence of UMI error correction (**Figure 2a, b**). Moreover, only UMI lengths of 8 or higher counted RNAs accurately over the full spectrum of assessed expression levels (**Figure 2c, d**).

**Figure 2:**
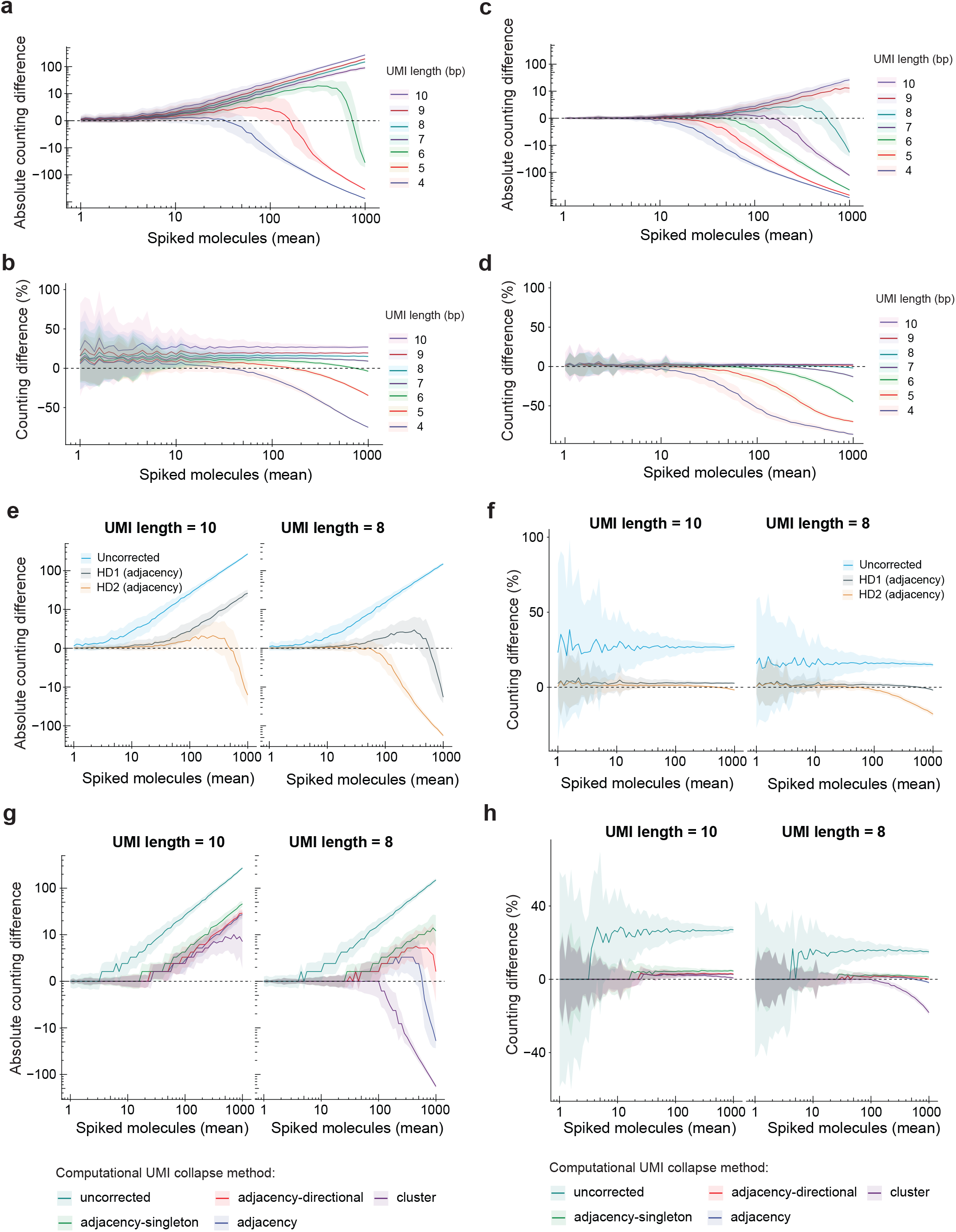
Evaluation of computational RNA counting strategies using molecular spikes. (**a-d**) Counting difference between number of unique spike identifiers and quantified 10x Genomics UMIs at varying mean expression levels. Colored lines indicate the mean (n=100) counting difference per UMI length shaded by the standard deviation. Counting difference is expressed in (a, c) absolute numbers or (b, d) as a percentage of the mean spUMI count and UMI counts were (a, b) computed without error-correction or (c, d) corrected in adjacency mode (hamming distance 1). (**e-f**) Comparison of edit distance (hamming distance) for adjacency error correction of UMIs of length 8 or 10. Lines indicate the mean (n=100) difference in quantification between spUMIs and UMIs shaded by the standard deviation in (e) absolute scale or (f) relative to the mean. (**g-h**) Evaluation of computational UMI collapse methods adjacency, adjacency-singleton, adjacency-direction and cluster at edit distance 1 and UMI lengths of 8 or 10 nucleotides. Lines indicate the mean (n=100) difference in quantification between spUMIs and UMIs shaded by the standard deviation in (g) absolute scale or (h) as a percentage relative to the mean.

Many common scRNA-seq pipelines have implemented UMI error corrections at an edit distance of 1, and we next compared the RNA counting accuracy by collapsing the same data using edit distances of 1 and 2 and compared the counts to the ground-truth based on the spiked in molecules. While a hamming distance of 1 was clearly more suitable for UMIs of length 8, allowing up to 2 mismatches in 10bp UMIs improved RNA counting throughout the full range of expression levels **(Figure 2e,f)**. Finally, we compared several UMI collapsing strategies that collapse UMIs based on their edit distances and frequencies of observations^7^ (**Supplementary Figure 6**) and we compared their inferred counts to the ground-truth spiked in molecules. Differences among the collapsing strategies were only apparent for UMIs of 8 basepairs in length (**Figure 2g,h**), where the aggressive collapsing strategies (*cluster* and *adjancency*) underestimate RNA counts due to the collapsing of multiple molecules at higher expression levels, likely due to coding space exhaustion. In line with previous findings^7^, the *directional-adjacency* method seems to provide a good compromise for UMIs of at least 8 base pairs.

RNA spike-in pools of different abundances and isoform complexities (e.g. ERCCs^13^, SIRVs^14^ and Sequins^15^) have been used to correlate known RNA molarities to observed RNA counts to assess gene- and isoform-level accuracy in scRNA-seq experiments^16^. Imprecisions in quantifying, diluting and pipetting minute amounts of spike-in mRNA to individual cells have however limited their general use and lowers their power to detect RNA counting errors or biases. Here, we propose molecular spikes, i.e. RNA spike-in pools that contain an inbuilt UMI (**Figure 1a,b**), as a new paradigm in scRNA-seq method development to detect and quantify artifactual RNA counting. Since the molecular spikes harbor an internal high-capacity UMI that can be used to quantitatively monitor the exact spike-in molecules sequenced from each cell, it is robust to imprecisions in accurately distributing spike-ins across cells. The quantitative comparison of spiked molecules to the counted RNA revealed both gross (e.g. 400%, **Figure 1g**) and smaller (5-10%) counting errors (**Figures 1e,g**), both relating to procedures that did not sufficiently remove UMI-containing oligonucleotides from contributing during PCR. We therefore suggest that molecular spikes should be routinely applied to existing and new scRNA-seq method development to validate accurate molecule counting, in particular when altering pre-PCR and PCR experimental conditions. To this end, we are making the molecular spikes available along with an R package for molecular spike data processing, quality-control, and visualization.

The generation of ground-truth molecular counts across cells with molecular spikes, enables systematic benchmarking of UMI error-correction strategies as one can quantitatively compare estimated RNA counts to the numbers of spiked molecules per cell. We show direct experimental evidence that RNA counting based on uncorrected UMIs over-estimate RNA expressions, at a level that follow the chance of PCR and sequencing errors within the UMIs (i.e. UMI lengths, sequence depth and sequencing technology used). In contrast to recent recommendations based on computational modelling^17^, our direct experimental comparison show that scRNA-seq data processing should include UMI error-correction to not systematically over-estimate RNA expression levels. The literature provides conflicting recommendations regarding UMI lengths^12,17^, as longer UMIs can interfere with method sensitivity and shorter UMIs have limited coding capacity. We demonstrate that only UMIs of 8 or more basepairs have sufficient coding capacity to robustly detect expression levels even in high RNA-content, cultured cells (here HEK293FT cells), and the use of shorter UMIs should be avoided except in shallow scRNA-seq experiments. Interestingly, none of the correction strategies typically used were fully robust across expression levels and it should be possible to use the quantitative data from the molecular spikes to inform future improved strategies with increasing RNA counting reliability and accuracy. It will also be interesting to use the molecular spikes beyond the validation of aggregated RNA counts per cell, and to investigate the within-molecule consistency of molecular spike identity and UMIs assigned to each molecule. In particular since an exact one-to-one mapping between sequence reads and original molecules (after the UMI error-correction) is important for in silico RNA reconstruction^6^ to ensure the correct collapsing of sequences for each individual RNA molecule present in cells.

## DATA AVAILABILITY

The raw data files for single-cell RNA-sequencing experiments have been deposited in Array Express at European Bioinformatics Institute under accession E-MTAB-10372.

## CODE AVAILABILITY

We are making the code for processing, filtering, quality control and visualization of molecular spikes publicly available as a R package (https://github.com/cziegenhain/UMIcountR).

## METHODS

### Molecular spike-in design

Molecular spike sequences were designed to have minimal overlap to Mouse or Human genomes. Two 500-bp sequences were selected, and entry vectors were created as described below. To minimize levels of *in vitro* transcription from the 5’ synthetic spike empty vector, we decided to complete the T7 promoter sequence with the random-base containing oligonucleotide. A similar strategy was not possible for the 3’ synthetic spike.

### 5’ and 3’ spike entry vector and library cloning

Geneblocks encoding synthetic RNA sequences and a synthetic poly-A stretch were introduced into the pUC19 backbone as previously described^6^. The resulting molecular spike insert vectors were linearized by digestion with XhoI or EcoRI for the 5’ and 3’ spike encoding plasmids respectively. A single stranded oligonucleotide library (IDT), containing a stretch of 18 random bases, was cloned into the linearized backbone using Gibson Assembly (NEB). The resulting reaction was then electroporated into Lucigen Electrocompetent cells, according to the manufacturers protocol, and streaked out on large LB-agar plates (LB-lennox recipe). The resulting cultures were recovered from the LB-agar plates and a maxipreps were performed (Macherey-Nagel) to purify the plasmid DNA.

### In vitro transcription reactions

The plasmid libraries were linearized by digesting with HindIII. In vitro transcription was performed using the MaxiScript kit (Invitrogen) according to the manufacturer’s guidelines. Resulting libraries of synthetic RNA spikes were cleaned up using RNeasy spin columns (Qiagen). Synthetic RNA integrity was confirmed by RNA Nano 6000 chip on the Agilent Bioanalyzer.

### Cell culture

HEK293FT cells (Invitrogen) were grown in complete DMEM medium supplemented with 4.5 g/L glucose, 6 mM L-glutamine, 0.1 mM MEM nonessential amino acids, 1 mM sodium pyruvate, 100 µg/mL pencillin-streptomycin and 10% fetal bovine serum (FBS). Prior to scRNA-seq experiments, cells were dissociated using TrypLE.

### 10x Genomics library preparation

1 ul of 3’ molecular spike pool (1 ng/µL) was added to the single cell HEK293FT suspension right before loading on the 10X genomics 3’ V2 chip. To avoid obtaining too many cells, and to remove the possibility of many ‘empty’ droplets that reverse-transcribed only the molecular spike molecules, we opted to remove GEMs from the reaction before the recovery of the cDNA. Before adding recovery agent, 10 µL of GEM-RT mix was transferred and the remainder of the GEM-RT mix was discarded. PCR amplification was performed according to the manufacturers protocol. After PCR amplification, cleanup was performed with SPRIselect beads at a ratio of 0.8:1 beads:sample instead of the 0.6:1 ratio specified in the protocol. The subsequent fragmentation step was extended to 10 minutes. The double-sided bead-cleanup after the fragmentation was changed to a ratio of 0.6:1 and 1:1 respectively. Similarly, the post-ligation cleanup (step 3.4) was increased to 1:1 ratio instead of 0.8:1. The double-sided post-indexing PCR cleanup was performed at 0.6:1 and 1:1 ratios respectively. The library was converted to circular ssDNA using the Universal Library Conversion Kit App-A (MGI). 60 fmol of ssDNA was used for DNA nanoball generation and subsequent sequencing on a FCL flow-cell of the DNBSEQ G400RS platform (MGI) generating 26×150 bp reads.

### Smart-seq3 library preparation

Single HEK293FT cells were sorted in 384 well plates containing 3 µL Smart-seq3 lysis buffer on a BD FACS Melody sorter with 100 µm nozzle. After sorting plates were quickly spun down before storage in -80 °C. Smart-seq3 library preparation was done according to published protocol^6^ with the following modifications. The 3uL Smart-seq3 lysis buffer per well contained 0.025 pg 5’ molecular spikes. After reverse transcription, each well containing 4 µL of cDNA was cleaned up with 3 µL home-made 22% PEG beads and eluted in 5 µL Tris-HCl pH 8. PCR mix was added as 5 µL to each well either with or without the addition of TSO. The reaction concentrations for the PCR in 10 µL were as follows: 1x KAPA HiFi Hot-Start PCR mix, 0.3 mM dNTPs/each, 0.5 mM MgCl2, 0 µM, 0.1 µM, 0.5 µM or 1.0 µM forward primer, 0.1 µM reverse primer. In the samples where TSO was added back into the PCR mix, it was done so at 0.8 µM.

### SCRB-seq library preparation

Single-cells were sorted into 96-wells containing 5 µL of lysis buffer (1/500 dilution of 5x Phusion HF Buffer) containing 0.025 pg of 3’ molecular spike pool using a BD FACSMelody sorter with 100 µm nozzle and frozen at -80 °C. After thawing, lysis was aided by Proteinase K digestion (1 µL of 1:20 diluted Proteinase K (Ambion)) for 15 min at 50 °C. Proteinase K was denatured, and RNA was desiccated by incubation at 95 °C for 10 min after unsealing the plate. Reverse transcription was performed in a volume of 2 µL per well (1 µM barcoded oligo-dT E3V6NEXT Biotin-ACACTCTTTCCCTACACGACGCTCTTCCGATCT[BC6][UMI10][T30]VN, 1x Maxima RT Buffer, 0.1 mM dNTPs, 1 µM TSO E5V6NEXT ACACTCTTTCCCTACACGACGCrGrGrG and 25 U Maxima H-reverse transcriptase) for 90 minutes at 42 °C. cDNA was pooled and cleaned using SPRI beads and excess primers digested by incubation with ExonucleaseI (NEB; 30 min @ 37 °C, inactivation 20 min @ 80 °C). PCR amplification was performed in 50 µL (0.5 µM SINGV6 primer Biotin-ACACTCTTTCCCTACACGACGC, 1x KAPA HiFi ReadyMix). PCR was cycled as follows: 3 min at 98 °C, 21 cycles of 15s at 98 °C, 30 s at 67 °C, 6 min at 72 °C and final elongation was performed for 10 min at 72 °C. For the direct PCR condition, we added 3 µL of PCR master mix directly to each well RT product well containing 2 µl of cDNA. Amplified, pooled cDNA was cleaned and quantified. 800 pg of cDNA was used for tagmentation using the Nextera XT kit (Illumina) according to the manufacturer’s protocol. The final indexing PCR was performed using a i7 primer and P5NEXTPT5 (AATGATACGGCGACCACCGAGATCTACACTCTTT CCCTACACGACGCTCTTCCG*A*T*C*T*; IDT) to select for correct 3’ fragments. The libraries were pooled and the converted to circular ssDNA using the Universal Library Conversion Kit App-A (MGI). 60 fmol of ssDNA was used for DNA nanoball generation and subsequent sequencing on a FCL flow-cell of the DNBSEQ G400RS platform (MGI) generating 16×150 bp reads.

### HEK293FT expression levels

UMI count tables from HEK293FT cells generated using the Smart-seq3 protocol were obtained from ArrayExpression accession E-MTAB-8735. After additional filtering of the cells (min. number of genes expressed 7,500 and minimum number of UMIs detected 50,000), we calculated the mean UMI count for all genes (n = 10,198) detected in at least 50% of the cells.

### Sequencing data processing

All sequencing data was processed using zUMIs (v2.9.5)^8^. Reads with more than 3 bases below Phred 20 base call scores in the UMI sequence were discarded. Remaining reads were mapped to the human genome hg38 and spike-in references using STAR (v2.7.3a)^18^ and mapped reads were quantified according to Ensembl gene models (Grch38.95) taking into consideration the strand information of the libraries. Error correction of the internal spUMI was applied within each cell barcode using the adjacency algorithm allowing edit distances of 2 (hamming distance).

### Computational analysis of molecular spikes

All downstream analyses were performed in R (v4.0.4). Reads aligning to the molecular spike reference sequence were loaded along with the library UMI and barcode information from zUMIs output bam files using Rsamtools^19^ and further processed by matching the known sequence upstream and downstream of the internal UMI. Only valid reads that had an 18 nucleotide long internal UMI were considered further.

To investigate the distances of uncorrected, hamming-distance 1 and hamming-distance 2 corrected spUMI sequences, we used the 5’-molecular spike data generated by the Smart-seq3 protocol and the 3’ molecular spike data from the 10x Genomics experiment. For each cell, we calculated all pairwise hamming distances of spUMI sequences within that cell as well as all pairwise distances to 1000 randomly sampled spUMI sequences across the whole dataset using the stringdist package^20^.

For the estimation of the complexity of the molecular spike pool, we counted the number of unique error-corrected spUMI sequences over molecules seen in all cells and fitted a non-linear asymptotic regression model using the NLSstAsymptotic function and extracted the asymptote (total complexity) from the coefficients of the model.

Overrepresented spike-ins were discarded if they were detected in more than 4 or 8 cells (5’- and 3’-spUMIs, respectively) or with more than 100 raw sequencing reads.

### Analysis of counting performance in protocol variations

For every cell barcode, spUMIs were randomly drawn from all molecular spike molecules within that barcode for 20 expression levels from 1 to 100 molecules. At each expression level and for each cell, we determined the exact number of molecules by drawing from a normal distribution with the given mean and added Poisson noise (standard deviation = square root of the mean). All observed sequencing reads associated to each of the drawn molecules were stored and adjacency error correction (hamming distance 1) was applied to the observed UMI sequences derived from the library preparation (for example the UMI in the Smart-seq3 TSO or the UMI in 10x Genomics oligo-dT).

### Evaluation of UMI length and UMI collapse algorithms

We first selected a pool of eligible spike-in molecules from all cells in the 10x Genomics dataset that fulfilled the following criteria (1) only observed in one cell barcode and (2) covered with 10 - 20 sequencing reads. From this pool of 26,815, we sampled molecules at 60 expression levels evenly spaced in log-space from 1 to 1000 molecules. At each expression level, we sampled for 100 “*in-silico*” cells the used number of spike molecules by drawing from a normal distribution with the given mean and added Poisson noise (standard deviation = square root of the mean). All associated sequencing reads were stored, and we shortened the UMI sequence in 1 base increments (3’ to 5’ direction) from 10 to 4 nucleotides. We then applied our R implementations of the following UMI error corrections within each expression level and in-silico cell: (1) *adjacency*: the network of closely related UMI sequences is resolved by collapsing all sequences within the given edit distance (ran with hamming distance 1 and 2 in our case) to the most abundant sequence; (2) *adjacency-directional*: as *adjacency*, but the minor nodes can only be collapsed if they have 0.5x or less the reads as the most abundant sequence. (3) *adjacency-singleton*: as *adjacency*, but the minor nodes can only be collapsed if they are observed by exactly 1 read; (4) *cluster*: the network of closely related UMI sequences is resolved by collapsing all sequences within the given edit distance to the node with the highest number of read counts. Nodes that were related at the same distance to one of the collapsed sequences and equally or less abundant are then also collapsed to the main node, even if their edit distance is higher than the initial parameter.

## ACKNOWLEDGEMENTS

We thank Henry and Urban Lendahl for valuable discussions and Matilda Eriksson from the Eukaryotic Single Cell Genomics facility (SciLifeLab, Stockholm) for help with the 10x Genomics library preparation. This work was supported by an EMBO long-term fellowship (ALTF 673–2017) grant to C.Z., an HFSP long-term fellowship (LT000155/2017-L) grant to G.-J.H, and grants to R.S. from the Swedish Research Council (2017-01062), the Knut and Alice Wallenberg Foundation (2017.0110), the Göran Gustafsson Foundation, and the Bert L. and N. Kuggie Vallee Foundation.

## AUTHOR CONTRIBUTION

Conceived the idea: GJ.H. Designed and cloned molecular spikes: GJ.H. Performed scRNA-seq experiments: C.Z., M.HJ., GJ.H. Performed analyses and generated figures: C.Z. Wrote the manuscript: R.S., C.Z. and GJ.H. Supervision: R.S.

## Supplementary Figure Legends

**Supplementary Figure 1:**
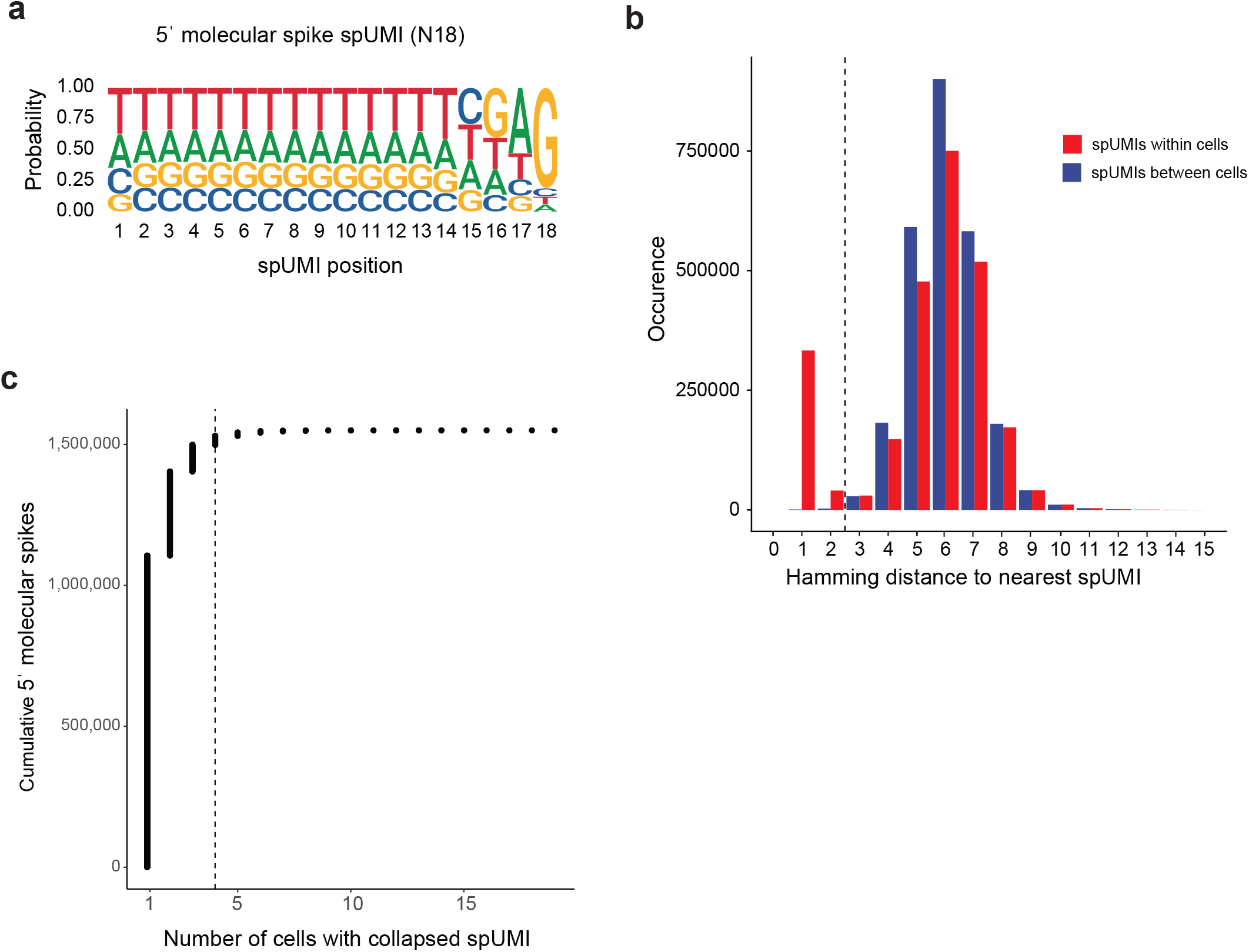
Quality control of 5’ molecular spike-in. (**a**) Sequence logo of the 18 random spUMI bases derived from all reads in the Smart-seq3 dataset. At each position, the frequency of all 4 bases is visualized by the size of the DNA letter. (**b**) Minimal distance of uncorrected spUMIs to the closest spUMI sequence for all pairwise within-cell comparisons and pairwise comparisons of spUMIs to 1000 randomly samples spUMI sequences across cells (total 2,233,878 comparisons). (**c**) Cumulative number of molecular spikes (n = 885,925) sorted by their occurrence over cells (n = 340). Dashed line indicates the chosen QC cutoff at 4 cells.

**Supplementary Figure 2:**
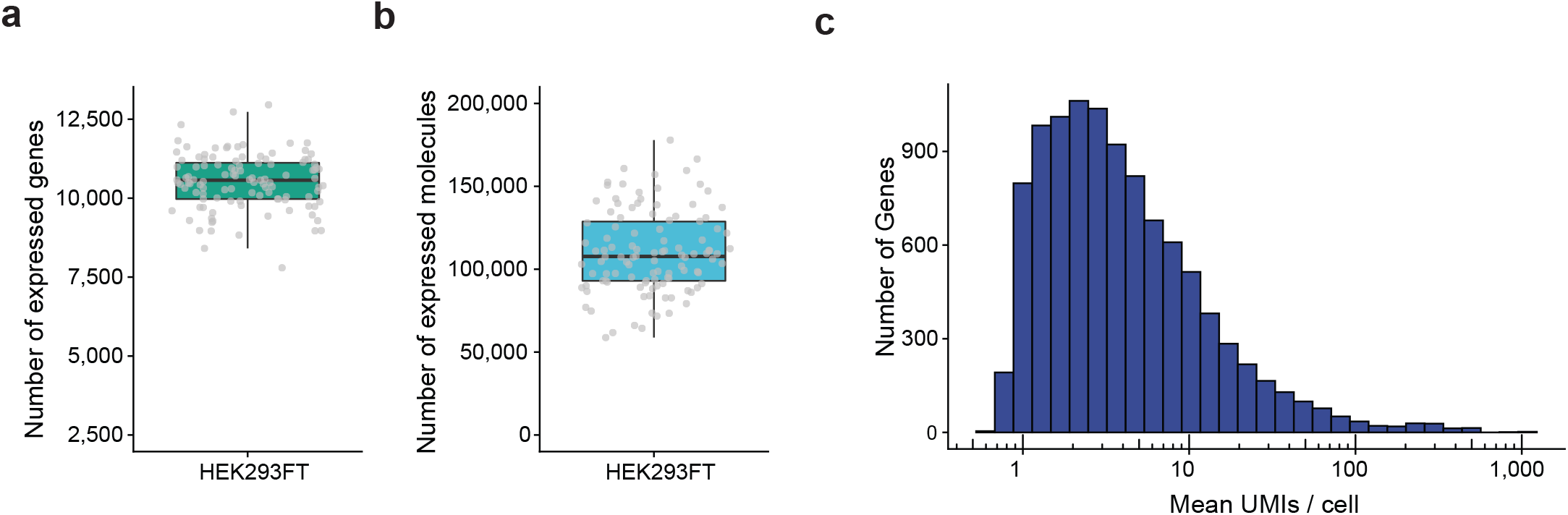
Expression levels in HEK293FT cells. (**a**-**b**) Quality of Smart-seq3 libraries (n = 111 cells) after filtering. Shown are the number of detected (a) genes and (b) molecules per HEK293FT cell. (**c**) Histogram showing the mean UMI count per cell for all genes expressed in at least 50% of cells (n = 10,198 genes).

**Supplementary Figure 3:**
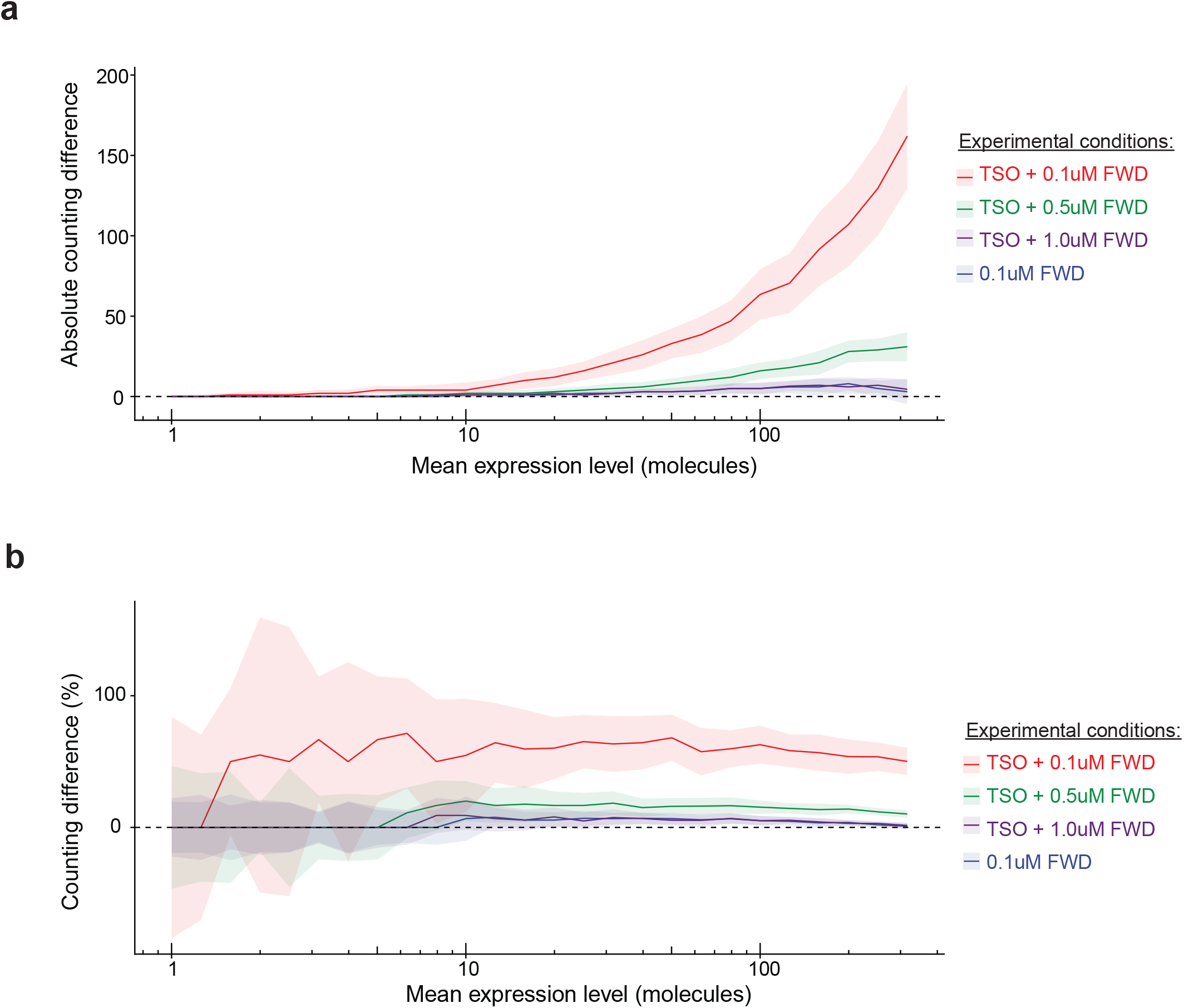
Counting difference in Smart-seq3 protocol variations. (**a**-**b**) For variations of the Smart-seq3 protocol, molecular spikes were sampled at varying mean expression levels. Colored lines indicate the mean counting difference in (a) absolute numbers or (b) relative to the mean and shaded by the standard deviation for library preparation conditions 0.1 µM FWD (n=48), TSO+0.1 µM FWD (n=48), TSO+0.5 µM FWD (n=39) and TSO+1.0 µM FWD (n=45).

**Supplementary Figure 4:**
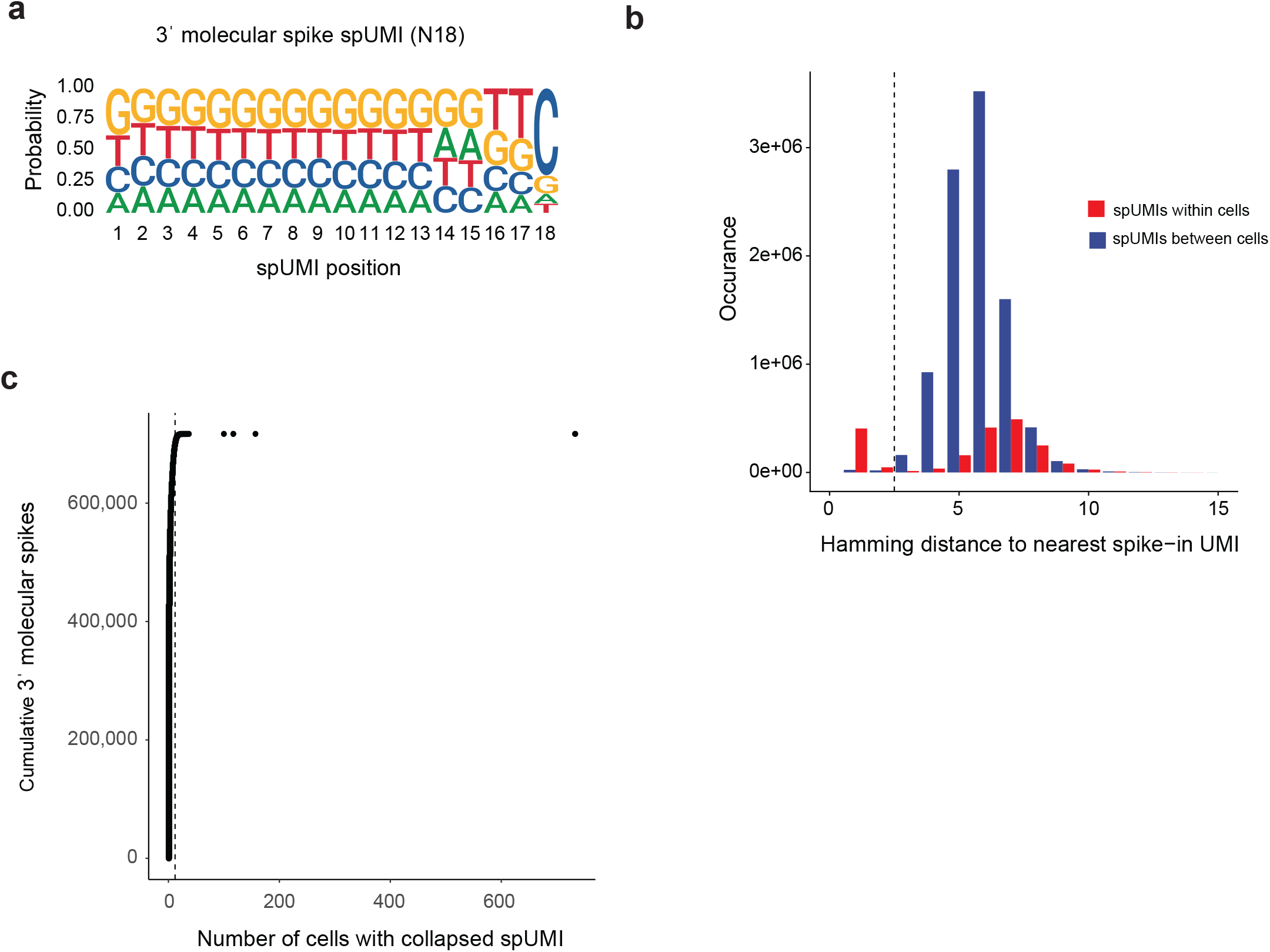
Quality control of 3’ molecular spike-in. (**a**) Sequence logo of the 18 random spUMI bases derived from all reads in the 10x Genomics dataset. At each position, the frequency of all 4 bases is visualized by the size of the DNA letter. (**b**) Minimal distance of uncorrected spUMIs to the closest spUMI sequence for all pairwise within-cell comparisons and pairwise comparisons of spUMIs to 1000 randomly samples spUMI sequences across cells (total 19,773,932 comparisons). (**c**) Cumulative number of molecular spikes (n = 1,938,392) sorted by their occurrence over cells (n = 1,359). Dashed line indicates the chosen QC cutoff at 4 cells.

**Supplementary Figure 5:**
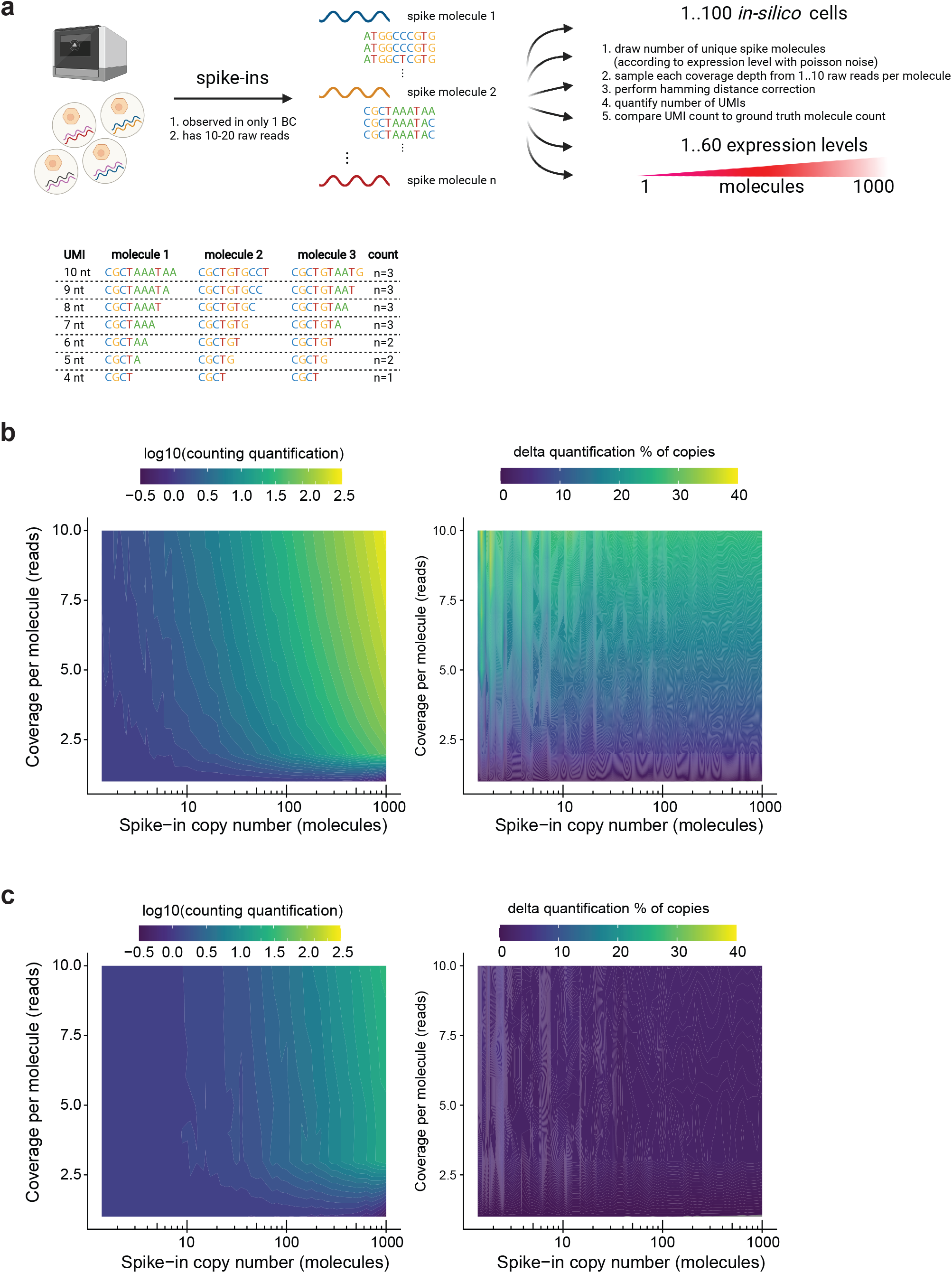
Strategy for sub-sampling molecular spikes to assess counting reliability across expression levels. (**a**) Strategy for computational analysis of 10x Genomics spUMI data. Molecular spike-ins observed in only one cell barcode and covered by 10 – 20 sequencing reads are selected along with their associated 10x UMI sequence. spUMIs were sampled at 60 expression levels ranging from 1 to 1000 molecules for 100 *in silico cells*. For each “cell” at each expression level, molecules were analyzed at depth of 1 to 10 reads and UMI error correction was applied. (**b**-**c**) We quantified the spUMIs and 10x UMIs and display the mean counting difference over the 100 replicates as a contour plot depending on expression level and read coverage in absolute numbers and normalized to the mean copy number, where (b) shows uncorrected 10x UMI counts and (c) shows UMI counts after applying an error correction at hamming distance 1.

**Supplementary Figure 6:**
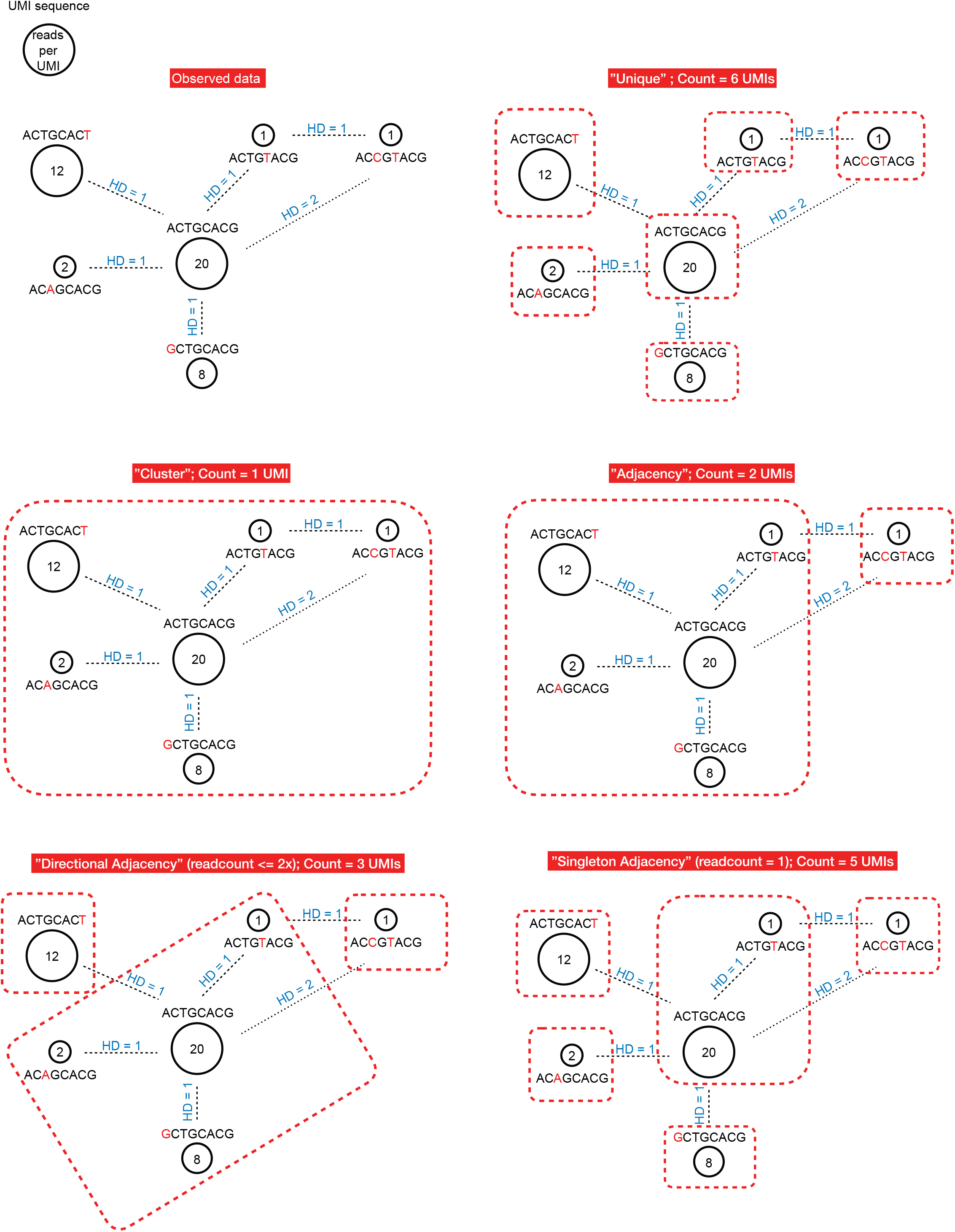
Computational algorithms for UMI collapsing. (**a**) Scenario of a network of UMI sequences where each UMI sequence is visualised along with the number of reads it was observed by. Mismatches to the center UMI sequence are shown in red and the edit distance (hamming distance HD) is indicated in blue. (**b**) Unique: Every unique sequence is counted as a molecule (naive counting, e.g. Kallisto). UMI count in the network = 6. (**c**) Cluster: The network is resolved by collapsing all sequences within HD1 to the UMI with the highest number of read counts. UMIs that were related at HD1 to one of the collapsed sequences and equally or less abundant are then also collapsed to the main UMI sequence, even if their edit distance is higher than 1. UMI count in the network = 1. (**d**) Adjacency: The network is resolved by collapsing all sequences within HD1 to the UMI with the highest number of read counts. UMI count in the network = 2. (**e**) Directional Adjacency: The network is resolved by collapsing all sequences within HD1 to the UMI with the highest number of read counts, unless they are observed with more than 50% of read support compared to the main UMI. UMI count in the network = 3. (**f**) Singleton Adjacency: The network is resolved by collapsing all sequences within HD1 and observed with only 1 read to the UMI with the highest number of read counts. UMI count in the network = 5.

